# Multivariate environmental exposures are reflected in whole-brain functional connectivity and cognition in youth

**DOI:** 10.1101/2025.11.13.688261

**Authors:** Sarah D. Lichenstein, Yihe Weng, Heather Robinson, Lester Rodriguez, Marzieh Babaeianjelodar, Joliza Maynard, Menessa Metayer, Suhani Suneja, Corey Horien, Abigail S. Greene, R. Todd Constable, Tyler M. Moore, Ran Barzilay, Sarah W. Yip, Arielle S. Keller

## Abstract

Each individual’s complex, multidimensional environment, known as their “exposome”, plays an essential role in shaping cognitive neurodevelopment. Understanding the mechanisms whereby children’s exposome influences their development is crucial to facilitate the design of interventions to foster positive developmental trajectories for all youth. Recent work has identified a general exposome factor associated with socio-economic inequality that is strongly related to cognition and individual differences in the spatial organization of functional brain networks in youth. Building on these findings, the current study explores whether alterations in functional connectivity may represent a potential mechanism linking variation in the exposome to cognitive performance. We apply a data-driven, cross-validated, whole-brain machine learning approach, connectome-based statistical inference, to identify patterns of functional connectivity associated with exposome scores among early adolescents enrolled in the Adolescent Brain Cognitive Development (ABCD) Study using data collected during three cognitive tasks and during rest. Additionally, we investigate whether the identified patterns of functional connectivity relate to individual differences in cognitive performance across three domains: General Cognition, Executive Functioning, and Learning/Memory. Models incorporating 10-fold cross-validation over 100 iterations identified consistent functional connections associated with the exposome across task and rest conditions (model performance: *ns* = 6,137 – 8,391, *r*s = 0.34 - 0.44, *p*s <.001). Results were robust across data collection sites and functional connections common across all significant models were associated with cognitive performance across domains (*p*s < 0.0009). Collectively, these findings reveal that multidimensional environmental exposures are reflected in patterns of functional connectivity and relate to cognitive functioning among youth.

**Highlights:**

- Patterns of functional connectivity reflect multivariate environmental exposures.
- Consistent exposome-related connectivity patterns were observed across brain states.
- The identified general exposome network was associated with cognitive performance across domains.

**Graphical Abstract:** 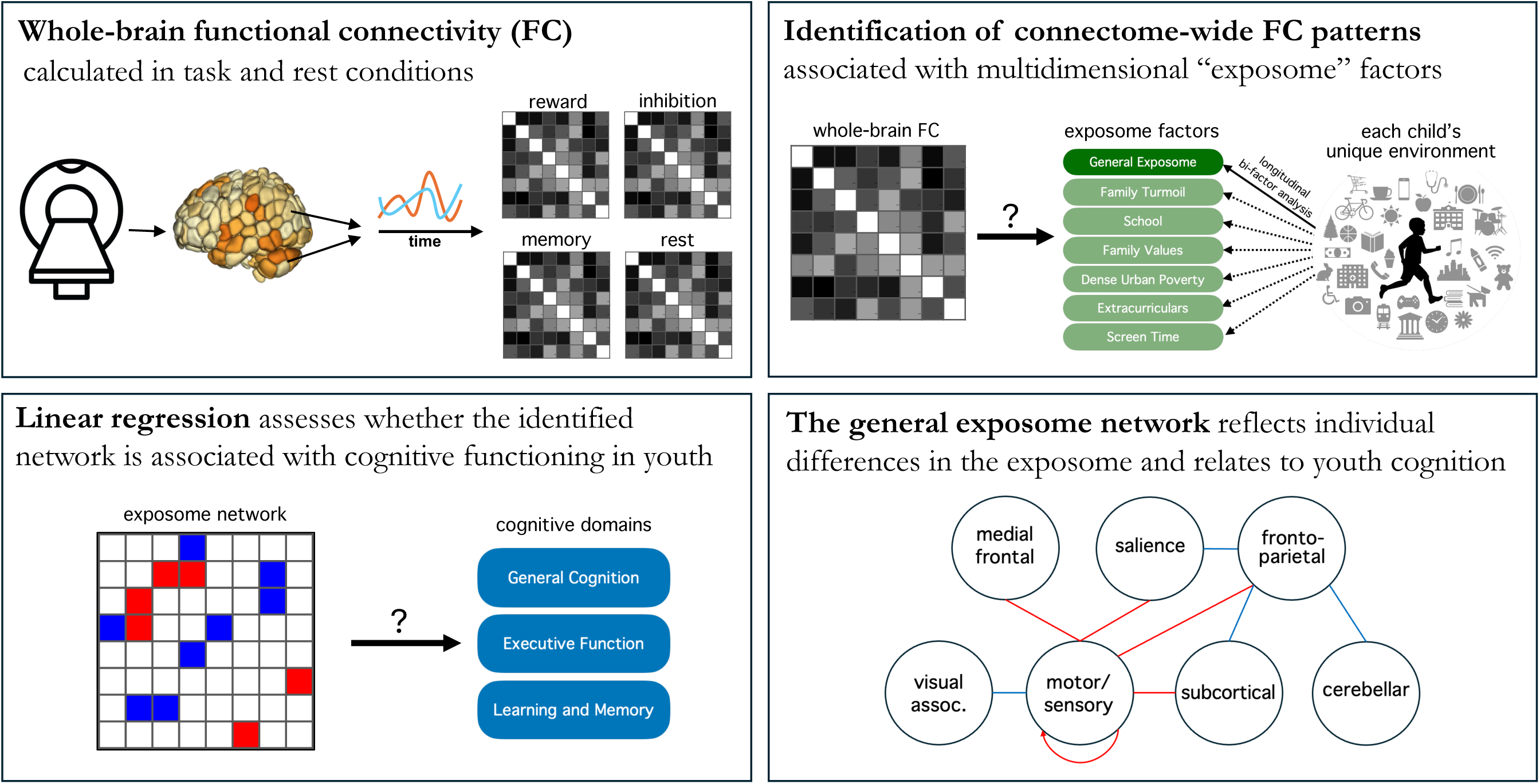

## 1. Introduction

Each individual’s unique environment and lived experiences play an essential role in shaping cognitive neurodevelopment. Previous work has identified numerous environmental factors that relate to neurodevelopment and cognition, including physical and chemical exposures (e.g. lead poisoning^1^ or air pollution^2,3^), psychosocial experiences (e.g. caregiver relationship^4,5^ or adverse life events^6^), and external environments (e.g. socioeconomic resources^7^ or cultural influences^8^). Most studies to date have examined associations with just one or a few environmental factors at a time. However, it is increasingly accepted that these features often co-occur non-randomly and likely have additive and interactive effects on the developing brain. For example, living in a high poverty neighborhood is associated with greater risk of exposure to air pollutants and traumatic experiences, which in turn are associated with individual differences in cognitive, psychiatric, and neurodevelopmental outcomes^2,9,10^. Consequently, there are increasing efforts to more comprehensively measure each individual’s complex, multidimensional environment, referred to collectively as an individual’s “exposome”. Efforts to characterize the exposome have begun to uncover important associations between large sets of co-occurring environmental features and individual differences in neurodevelopmental outcomes, with potential to ultimately inform interventions that support healthy neurocognitive development^11^.

The concept of the “exposome” was first introduced as a complement to the genome in epidemiological studies^12^, emphasizing the importance of capturing the totality of co-occurring environmental exposures in addition to genetic influences on physical health. Recently, this exposome approach has been embraced by a wider range of scientific fields and definitions of the exposome have expanded to include psychosocial and sociocultural factors^13^. Emerging work has already begun to leverage large-scale datasets, such as the Adolescent Brain Cognitive Development℠ Study (ABCD StudyⓇ)^14^, to investigate associations between the exposome and brain network organization^15^, cognitive outcomes^15–17^, and mental health^17,18^. For example, a multidimensional exposome score reflecting lower socioeconomic status (SES) is associated with poorer cognitive performance and worse mental and physical health^17^. In addition, including multidimensional environmental exposures improves brain-based predictions of cognitive performance^19^ beyond what can be gained by including single environmental features such as household income or neighborhood deprivation indices. These approaches have leveraged bifactor analysis to quantify both a single general exposome factor and several sub-factors that capture specific dimensions of the enrivonment^15,18^. The general exposome factor uncovered using this bifactor approach in the ABCD Study captures co-occurring environmental features associated with socio-economic inequality and is strongly associated with symptoms of psychopathology^18^, cognitive performance^15^, and individual differences in the spatial organization of functional brain networks^15^ in youth.

Building on these findings, the current study aims to explore whether alterations in functional connectivity may represent a potential mechanism linking variation in the exposome to cognitive performance. Coordinated patterns of activity across distributed networks of functionally related brain regions have been shown to give rise to complex cognitive functions^20^, and individual differences in whole-brain functional connectivity profiles have predicted cognitive performance across multiple domains, including fluid intelligence^21,22^, attention^23,24^, language^25^ and executive functioning^25^. Moreover, developmental changes in functional connectivity profiles^26–29^ are associated with age-related improvements in cognitive performance^30–33^, suggesting that functional connectivity profiles may be a promising, developmentally sensitive biomarker of individual differences in youth cognition. Recent work has also shown that functional connectivity relates to environmental exposures, including household socioeconomic status^34,35^, neighborhood disadvantage^34^, interpersonal unpredictability^35^, and air pollution^36^, with important implications for cognition and mental health^35,37,38^. However, prior work has focused on examining only a select subset of environmental features at a time, and the relationship between the full exposome and whole-brain functional connectivity patterns during development remains unknown.

Here, we apply a modified version of a well-validated, data-driven, whole-brain, machine learning approach, connectome-based predictive modeling (CPM), which we here refer to as connectome-based statistical inference (details below), to identify patterns of functional connectivity associated with exposome scores among early adolescents enrolled in the ABCD Study. Considering recent evidence that different task conditions, i.e. “brain states”, are optimal for revealing distinct brain-phenotype relationships with CPM and other predictive modeling approaches^22,39–42^, we independently examine patterns of functional connectivity using functional magnetic resonance imaging (fMRI) data collected during three different cognitive tasks and during rest. Additionally, we investigate whether the identified patterns of functional connectivity associated with each individual’s exposome also relate to individual differences in cognitive performance across three domains: General Cognition, Executive Functioning, and Learning/Memory^43^. Collectively, this work aims to elucidate how multivariate childhood exposures relate to whole-brain functional connectivity and cognition among youth^15^.

## 2. Methods

### 2.1. Participants

Data were drawn from the Adolescent Brain Cognitive Development (ABCD) Study (*n*=11,878), a large-scale study of adolescent neurodevelopment with youth recruited from 21 sites across the United States at age 9-10 and followed prospectively across adolescence^14^. The study includes comprehensive demographic, cognitive, and clinical assessments, as well as neuroimaging, including fMRI during 3 tasks and rest (see 2.2.2., below). Details on the recruitment strategy have been previously described^44^. All study procedures were approved by a central Institutional Review Board (IRB) at the University of California, San Diego, and several sites also obtained local IRB approval. A parent or guardian provided written informed consent and assent to participate was obtained from participants.

### 2.2. Measures

#### 2.2.1. Exposome

We leveraged a set of previously derived exposome factors, including a general (overall) factor and 6 subfactor scores, defined using longitudinal bifactor analysis as previously described^15^. Briefly, general and specific exposome factors were derived from a set of 354 variables comprising youth-report, caregiver-report, and geo-coded data covering a wide range of environmental features, which were reduced to a set of 32 data-driven summary scores prior to exploratory structural equation modeling with bifactor rotation (for details see Keller *et al*.^14^). To ensure consistency with prior work, we used the exact same scores as in our previously published study rather than recalculate exposome scores within the present sub-samples of ABCD data across each cross-validation fold, noting that the derivation of the exposome factors was agnostic to measures of neural or cognitive function. Measures of socioeconomic status (SES) (e.g., household income, parent education) show the strongest loadings for the general exposome factor, whereas the 6 subfactors capture more specific dimensions of a child’s environment, each of which is orthogonal to the general exposome factor and all other subfactors. These subfactors include School (i.e., school involvement, enjoyment, and performance), Family Values (i.e., Mexican American Cultural Values Scale subscales measuring family values, centrality, and culture), Family Turmoil (i.e., family conflict based on both youth and parent reports), Dense Urban Poverty (i.e., geocoded and parent-report data on youth’s neighborhood, including poverty, density, safety, and pollution), Extracurriculars (i.e., involvement in sports and other activities and traumatic brain injury), and Screen Time.

#### 2.2.2. Neuroimaging Data

ABCD participants completed a neuroimaging assessment at baseline, including fMRI scanning using 3T scanners during rest and 3 cognitive tasks: the Stop-Signal Task (SST; response inhibition), the Monetary Incentive Delay Task (MID; reward processing), and the Emotional N-Back Task (EN-back; working memory, affective processing). Details regarding acquisition parameters and task design have been previously described^45^. Raw time series for the baseline fMRI data were obtained via ABCD’s Fast Track Release of raw fMRI data and preprocessed using the Yale Magnetic Resonance Research Center (MRRC) functional connectivity pipeline to generate functional connectivity matrices for each individual for each task and resting-state scan.

Preprocessing was conducted using Bioimage Suite^46^ and SPM12 including the entire raw time series, consistent with prior CPM work^42,47–50^, and included brain extraction^51^, nonlinear registration to MNI space, and motion correction. The following covariates were regressed out of the data: linear, quadratic, and cubic drifts, mean global signal, mean cerebral-spinal-fluid, mean white-matter signal, and a 24-parameter motion model including six rigid-body motion parameters, six temporal derivatives, and these terms squared. Participants with mean framewise displacement (FD) > 0.2 mm across runs of a given task were excluded from the analysis^52^ and variance attributable to mean FD was also accounted for in our primary analyses (see 2.3.1., below). Following preprocessing and exclusion for motion exceeding the 0.2 mm mean FD threshold, there were n=8,730 participants with functional connectivity and exposome data that were included across our primary analyses (see Table 1 for demographics; Rest: n=9,956 with successfully preprocessed functional connectivity data, i.e., good quality skull stripping and registration, n=8,877 following exclusion for head motion > 0.2 mm mean FD, n=8,391 with usable functional connectivity, exposome, sex, and age data for inclusion in analysis; MID: n=8,924 after preprocessing, n=6,936 after motion exclusion, n=6,672 with complete data for analysis; SST: n=8,827 after preprocessing, n=6,592 after motion exclusion, n=6,385 with complete data for analysis; EN-back: n=8,755 after preprocessing, n=6,334 after motion exclusion, n=6,137 with complete data for analysis).

**Table 1.** Demographic characteristics of participants included in primary analyses (n=8730)

| Variable | *M* | *SD* |
| --- | --- | --- |
| **Age (months)** | 119.20 | 7.52 |
| **Sex** | N | % |
| Female | 4431 | 50.76 |
| Male | 4299 | 49.24 |
| **Household Income** | N | % |
| <50K | 2164 | 24.79 |
| >=50K & <100K | 2337 | 26.78 |
| >=100K | 3524 | 40.38 |
| **Race** | N | % |
| White | 4817 | 55.18 |
| Black | 1154 | 13.22 |
| Hispanic | 1670 | 19.13 |
| Asian | 175 | 2.00 |
| Other | 913 | 10.46 |
| **Parent Education** | N | % |
| High school or lower | 1362 | 15.60 |
| Undergraduate or associates | 5069 | 58.06 |
| Advanced degree | 2290 | 26.23 |

Functional data were parcellated into 268 nodes using the Shen 268 atlas^53^, which includes cortical and subcortical regions, as well as the cerebellum. Functional connectivity between each node pair was then computed using Pearson correlations and correlation coefficients were transformed using the Fisher’s r-to-z transformation to yield 268x268 connectivity matrices, termed ‘connectomes’. These connectomes represent each participant’s whole-brain functional connectivity during each task and rest, which served as the primary input for our primary analyses (described in 2.3.1., below).

#### 2.2.3. Cognition

Cognitive performance was assessed using 3 principal components that have been previously derived from the ABCD neurocognitive battery using Bayesian Probabilistic Principal Components Analysis^43^, including components reflecting General Cognition, Executive Functioning, and Learning/Memory. These components were computed from seven measures from the NIH Toolbox, as well as the Rey auditory verbal learning test and the Little man task. Scores were obtained via the ABCD Data Exploration and Analysis Portal (DEAP).

### 2.3. Analysis

#### 2.3.1. Exposome Network Identification

Here we adapted a cross-validated machine learning approach, connectome-based predictive modeling (CPM), to identify neural networks that are statistically associated with individual differences in the exposome^42,47–50^ (hereafter referred to as connectome-based statistical inference). We use this term to recognize that the exposome was not calculated separately for individual folds, as described above. We note that while the exposome is treated as the outcome variable in these analyses, consistent with the CPM-style framework, the current application of connectome-based statistical inference aims to identify patterns of functional connectivity associated with individual differences in the exposome without implying any causality or directionality between brain and environmental variation. Individual participant connectomes computed from MID, SST, EN-back, and resting-state data served as the input for our primary analyses. We conducted 10-fold cross-validation, where 90% of the sample was used as training data and 10% was used as testing data within each fold. For the initial feature selection step, linear regression was used to identify edges that are significantly related to the outcome at *p* < 0.05 in the training data. To account for effects of sex, age, and head motion, these variables were regressed out of the exposome scores within each fold and the residuals of this analysis were used as the primary outcome measure. Edges for which stronger connectivity was a positive predictor of the residualized exposome scores and those for which reduced connectivity was a positive predictor of the residualized exposome scores were summed separately to generate positive and negative summary scores, respectively.

This was followed by a model generation step, in which a linear model was fit to predict the residualized exposome scores from the positive and negative summary scores. This model was then applied to the testing data to generate out-of-sample predictions of residualized exposome scores. The covariates in the test set were entered into the covariate regression model generated from the training set to yield predicted values reflecting the contribution of confounding variables. The final predicted exposome score was computed as the sum of the predicted residualized exposome score and the predicted covariate effects. Finally, model performance was evaluated based on the Pearson correlation between predicted and actual exposome scores across all participants in the test set.

To ensure that the model was not overfit to one 10-fold split, we repeated the analysis described above 100 times, using random splits of the data on each iteration, and calculated mean performance across all iterations. Because analyses across folds are not entirely independent, we tested for statistical significance via permutation testing, where the correspondence between connectomes and exposome scores was randomly shuffled and the model was rerun 1000 times to create a null distribution. Models were run separately for each task and rest, as well as separately for the general exposome factor and the 6 subfactors. To account for multiple comparisons across four brain states and seven exposome factors, a Bonferroni-corrected threshold of *p* < 0.002 was used to evaluate model significance. We then visualized the anatomy of identified networks based on overlap with canonical neural networks^54^ and anatomical location of high-degree nodes (i.e., nodes with the greatest number of significant edges), as in prior CPM work^42,47–50^. Cosine similarity was used to quantify similarity between networks identified across task conditions.

#### 2.3.2. Sensitivity Analyses

To assess variation in model performance across ABCD data collection sites, follow-up analyses were conducted quantifying the association between predicted and actual general exposome scores in each brain state across all participants from each site independently. To determine whether model performance varied systematically across data collection sites based on site differences in exposome variability, we then compared these association values with the standard deviation of exposome scores at each site.

#### 2.3.3. Associations with Cognition

To investigate whether the patterns of functional connectivity associated with exposome scores also relate to individual differences in cognition, we performed linear regression analyses to quantify the relationship between network strength of a consensus network (i.e., the network corresponding to edges consistently identified across brain states, details in Results) and each of the three cognitive scores (i.e., General Cognition, Executive Function, Learning/Memory), controlling for age and sex (head motion was not included as a covariate because it was accounted for during network identification). Significance was assessed using permutation testing, where the correspondence between network strength and each cognitive domain was shuffled 10,000 times to create a null distribution.

## 3. Results

### 3.1. Exposome network identification

#### 3.1.1. Model performance

Models incorporating data collected during all tasks and rest were successful in predicting the general exposome factor (*r*s = 0.34 - 0.44, *p*s < .001) and each of the 6 subfactors (*r*s = 0.08 - 0.25, *p*s < .001; see Table 2 for individual model performance statistics) in held-out participants.

**Table 2.** Model Performance.

| Brain State | Network | General Exposome |  |  | School |  |  | Family Values |  |  | Family Turmoil |  |  | Dense Urban Poverty |  |  | Extracurriculars |  |  | Screen Time |  |  |
| --- | --- | --- | --- | --- | --- | --- | --- | --- | --- | --- | --- | --- | --- | --- | --- | --- | --- | --- | --- | --- | --- | --- |
|  |  | *R* | *R*<sup>2</sup> (%) | *P* | *R* | *R*<sup>2</sup> (%) | *P* | *R* | *R*<sup>2</sup> (%) | *P* | *R* | *R*<sup>2</sup> (%) | *P* | *R* | *R*<sup>2</sup> (%) | *P* | *R* | *R*<sup>2</sup> (%) | *P* | *R* | *R*<sup>2</sup> (%) | *P* |
| Rest | Positive | 0.39 | 15.56 | <0.001 | 0.18 | 3.12 | <0.001 | 0.09 | 0.82 | <0.001 | 0.09 | 0.88 | <0.001 | 0.25 | 6.36 | <0.001 | 0.10 | 0.97 | <0.001 | 0.17 | 2.88 | <0.001 |
| Rest | Negative | 0.45 | 20.60 | <0.001 | 0.18 | 3.09 | <0.001 | 0.08 | 0.71 | <0.001 | 0.08 | 0.71 | <0.001 | 0.23 | 5.46 | <0.001 | 0.10 | 1.08 | <0.001 | 0.16 | 2.66 | <0.001 |
| Rest | Combined | 0.44 | 18.96 | <0.001 | 0.18 | 3.12 | <0.001 | 0.09 | 0.82 | <0.001 | 0.09 | 0.87 | <0.001 | 0.24 | 6.00 | <0.001 | 0.10 | 1.10 | <0.001 | 0.17 | 2.84 | <0.001 |
| SST | Positive | 0.32 | 10.24 | <0.001 | 0.16 | 2.70 | <0.001 | 0.08 | 0.61 | <0.001 | 0.09 | 0.86 | <0.001 | 0.23 | 5.18 | <0.001 | 0.08 | 0.71 | <0.001 | 0.12 | 1.55 | <0.001 |
| SST | Negative | 0.35 | 12.51 | <0.001 | 0.16 | 2.59 | <0.001 | 0.07 | 0.44 | <0.001 | 0.10 | 1.01 | <0.001 | 0.22 | 4.78 | <0.001 | 0.09 | 0.84 | <0.001 | 0.13 | 1.70 | <0.001 |
| SST | Combined | 0.34 | 11.81 | <0.001 | 0.16 | 2.67 | <0.001 | 0.08 | 0.57 | <0.001 | 0.10 | 0.97 | <0.001 | 0.23 | 5.08 | <0.001 | 0.09 | 0.83 | <0.001 | 0.13 | 1.68 | <0.001 |
| MID | Positive | 0.32 | 10.08 | <0.001 | 0.15 | 2.23 | <0.001 | 0.09 | 0.84 | <0.001 | 0.08 | 0.62 | <0.001 | 0.23 | 5.45 | <0.001 | 0.10 | 1.07 | <0.001 | 0.12 | 1.39 | <0.001 |
| MID | Negative | 0.36 | 13.14 | <0.001 | 0.15 | 2.21 | <0.001 | 0.09 | 0.88 | <0.001 | 0.08 | 0.67 | <0.001 | 0.22 | 4.65 | <0.001 | 0.10 | 1.01 | <0.001 | 0.12 | 1.41 | <0.001 |
| MID | Combined | 0.35 | 12.08 | <0.001 | 0.15 | 2.21 | <0.001 | 0.10 | 0.94 | <0.001 | 0.08 | 0.68 | <0.001 | 0.22 | 5.06 | <0.001 | 0.11 | 1.13 | <0.001 | 0.12 | 1.43 | <0.001 |
| EN-back | Positive | 0.37 | 13.65 | <0.001 | 0.18 | 3.39 | <0.001 | 0.10 | 1.10 | <0.001 | 0.10 | 0.98 | <0.001 | 0.24 | 5.99 | <0.001 | 0.10 | 1.01 | <0.001 | 0.13 | 1.73 | <0.001 |
| EN-back | Negative | 0.40 | 16.39 | <0.001 | 0.18 | 3.09 | <0.001 | 0.10 | 1.05 | <0.001 | 0.10 | 1.05 | <0.001 | 0.22 | 5.03 | <0.001 | 0.10 | 0.99 | <0.001 | 0.13 | 1.68 | <0.001 |
| EN-back | Combined | 0.40 | 15.84 | <0.001 | 0.18 | 3.28 | <0.001 | 0.11 | 1.20 | <0.001 | 0.10 | 1.03 | <0.001 | 0.23 | 5.52 | <0.001 | 0.10 | 1.07 | <0.001 | 0.13 | 1.77 | <0.001 |

Note. $R^2 = 1 - (\text{error SS})/(\text{total SS})$ . Abbreviations: SST: Stop-Signal Task; MID: monetary incentive delay task; EN-back: Emotional N-back Task.

#### 3.1.2. Network Anatomy

Consistent with prior CPM work, identified networks are complex and include edges connecting regions throughout the brain. Notably, consistent patterns emerged across general exposome networks identified using data from each task and rest (cosine similarity based on canonical functional network edge counts > 0.9, *p*s < 0.001; see Table 3). Therefore, we focus primarily on a consensus network composed of edges present in all identified networks, including 2,211 positive network edges and 1,839 negative network edges (see Figure 1; see Figure S1 for anatomy of networks generated from each task and rest condition separately).

**Figure 1.**
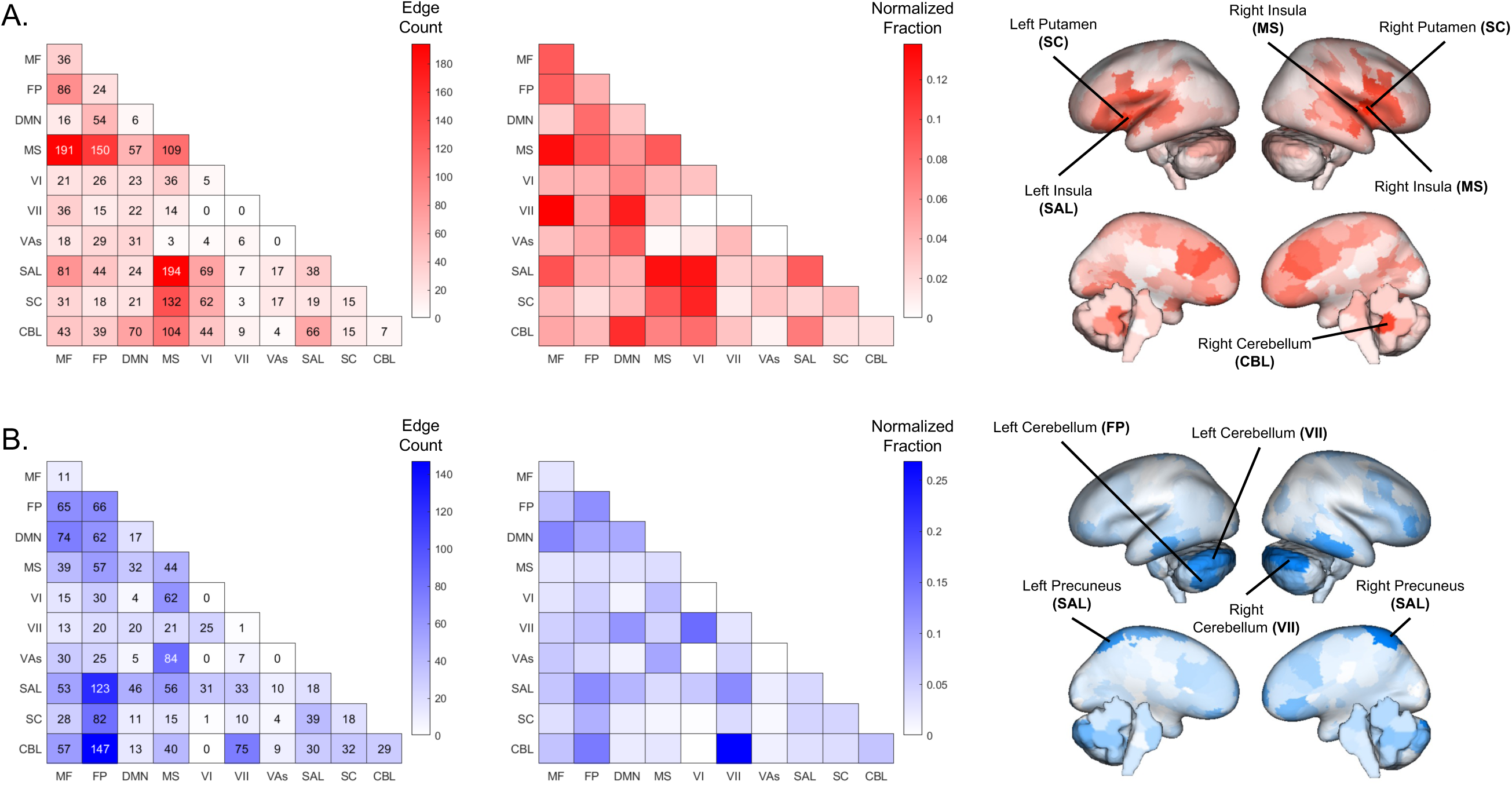
Consensus General Exposome Network Anatomy. Anatomy of edges identified in models conducted across all brain states (i.e., rest, MID, SST, EN-back) visualized based on overlap with canonical neural networks and nodal degree. Overlap with canonical neural networks is presented based on raw edge count and normalized by network size based on the total number of possible within/between-network edges (i.e., normalized fraction). Panel A shows positive network edges (n=2,211), for which greater connectivity is predictive of higher general exposome scores; panel B shows negative network edges (n=1,839), for which reduced connectivity is predictive of higher general exposome scores. MF: medial frontal, FP: frontoparietal, DMN: default mode network, MS: motorsensory, VI: visual 1, VII: visual 2, VAs: visual association, SAL: salience, SC: subcortical, CBL: cerebellar.

**Table 3.** Cosine similarity between general exposome networks identified across brain states.

|  | **SST** | **MID** | **EN-back** |
| --- | --- | --- | --- |
| **MID** | 0.953 |  |  |
| **EN-back** | 0.954 | 0.940 |  |
| **rest** | 0.953 | 0.902 | 0.935 |

Examining network anatomy based on overlap with canonical neural networks (Fig. 1A), higher general exposome scores, which reflect higher SES, residing in safer and less crowded neighborhoods, and having greater involvement in extracurricular activities, were associated with greater connectivity between the motorsensory network and medial frontal, frontoparietal, salience, and subcortical networks. Higher general exposome scores were also associated with lower connectivity between the frontoparietal network and salience, subcortical, and cerebellar networks, as well as between motorsensory and visual association networks. Examining network anatomy based on nodes with the most significant edges, i.e., high degree nodes (Fig. 1B), the highest degree nodes of the positive network included bilateral putamen, bilateral insula, and bilateral cerebellum. The highest degree nodes of the negative network included bilateral precuneus and bilateral cerebellar regions.

### 3.2. Sensitivity Analyses

To assess whether relationships between the identified networks and exposome scores vary across data collection sites, we conducted post-hoc sensitivity analyses by assessing the correlation between predicted and actual exposome scores within each site individually. Model performance remained significant across all sites for rest, and most sites across tasks (95.2% of sites for EN-back and MID, 85.7% of sites for SST), suggesting that our primary models are robust to site variation. Nonetheless, the strength of the association varied (see Figure 2). Notably, across all tasks and rest, model performance (i.e., the relationship between predicted and actual general exposome scores) was relatively stronger within the LA Children’s Hospital, South Carolina, New York, University of Florida, Maryland, Michigan, Missouri, and Connecticut sites, and relatively weaker within the Colorado, Florida International, Oregon, Minnesota, Utah, and Vermont sites, compared to model performance in the full dataset. This pattern appears to be at least partially attributable to greater variation in exposome scores across participants at the sites characterized by better model performance (see Figure S2).

**Figure 2.**
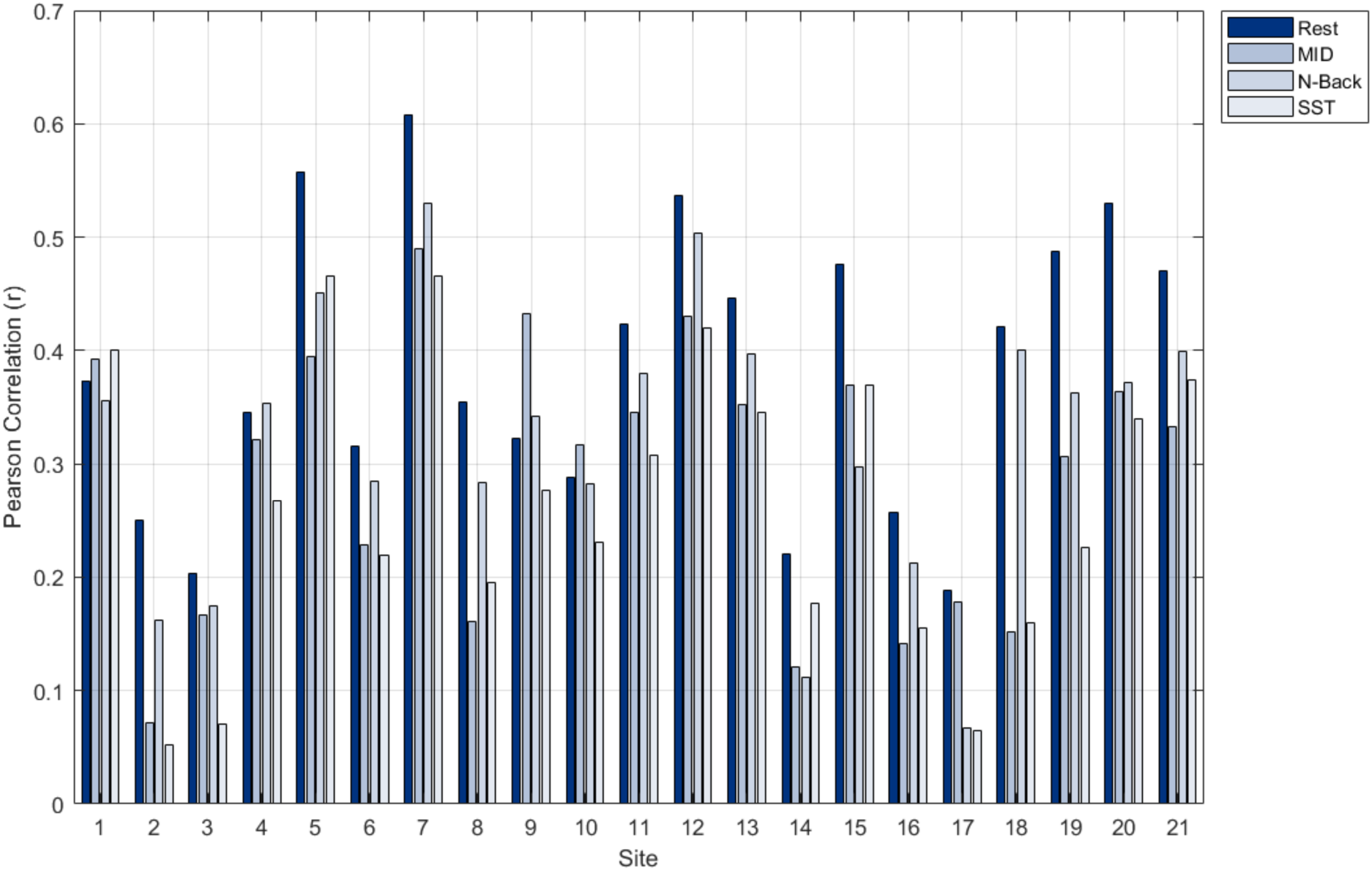
Variation in General Exposome Model Performance Across Sites. The association between model predicted and observed general exposome scores is presented within each site separately for models run on data collected during rest, MID, EN-back, and SST task performance. Model performance was significant across all sites based on permutation testing.

### 3.3. Associations with cognition

Greater consensus network strength was significantly positively associated with General Cognition (β*s* = 0.15-0.17), Executive Function (β*s* = 0.03-0.07), and Learning/Memory (β*’s* = 0.06-0.09; see Table 4 for individual model statistics and Figure 3).

**Figure 3.**
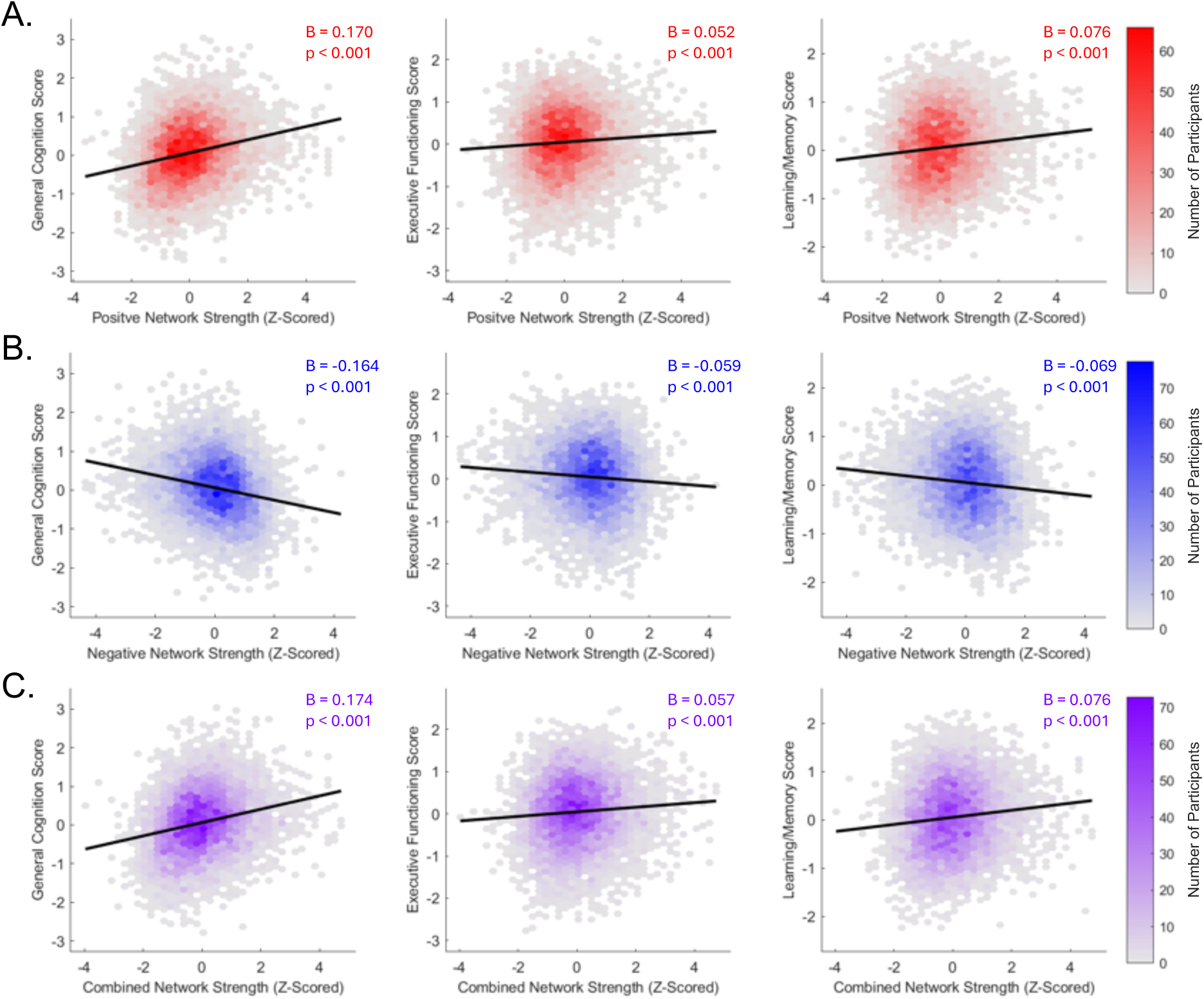
Associations Between Consensus General Exposome Network Strength and Cognition. The positive (A), negative (B), and combined (C) consensus general exposome network is significantly associated with General Cognition, Executive Function, and Learning/Memory during rest.

**Table 4.** Associations between General Exposome Combined Network Strength and Cognition across Brain States.

| Brain State | General Cognition |  | Executive Functioning |  | Learning/Memory |  |
| --- | --- | --- | --- | --- | --- | --- |
| | $\beta$ | Permuted Sig. | $\beta$ | Permuted Sig. | $\beta$ | Permuted Sig. |
| Rest | 0.17 | < 0.0001 | 0.06 | < 0.0001 | 0.08 | < 0.0001 |
| EN-back | 0.15 | < 0.0001 | 0.04 | < 0.0001 | 0.07 | < 0.0001 |
| SST | 0.15 | < 0.0001 | 0.03 | 0.0008 | 0.06 | < 0.0001 |
| MID | 0.16 | < 0.0001 | 0.07 | < 0.0001 | 0.09 | < 0.0001 |

## Discussion

Individuals’ early environments vary in myriad ways, with important implications for neurodevelopment. The current study applied connectome-based statistical inference – an adapted version of a well-validated, whole-brain, statistical approach, connectome-based predictive modeling – to identify patterns of functional connectivity predictive of individual differences in a general exposome factor capturing variation in over 350 variables reflecting different aspects of the environment. Consistent neural features were identified across models using data from three different cognitive tasks and rest. Furthermore, this general exposome network was robust to data collection site and predictive of cognitive performance across three domains: General Cognition, Executive Functioning, and Learning/Memory. Collectively, these findings provide novel insight into how multidimensional environmental exposures are reflected in patterns of functional connectivity and relate to cognitive functioning among youth.

Overall, early environments characterized by higher socioeconomic status were associated with greater connectivity among motorsensory, frontoparietal, medial frontal, salience, and subcortical network regions, coupled with reduced connectivity between frontoparietal, salience, cerebellar, and visual network regions. This pattern of results is consistent with prior ABCD studies demonstrating altered motorsensory^19,55^, frontoparietal^55^, salience^55^, cerebellar^19^, and subcortical^19^ connectivity in association with neighborhood deprivation^19,55^ and differences in the family environment^19^ (i.e., parent income and education), as well as findings from other studies reporting associations between air pollution and connectivity of motorsensory and salience networks^36^. These results also align with findings from prior ABCD work applying predictive modelling to elucidate functional connectivity-based substrates of cognitive performance, including greater motorsensory connectivity with medial frontal and subcortical regions and lower connectivity between frontoparietal and salience networks^56^. The central role for frontoparietal, motorsensory, visual and subcortical network connectivity is also consistent with prior work in other datasets applying CPM to predict various domains of cognitive performance^22,25,57^.

The current pattern of results also aligns with prior work suggesting that hierarchical development along a sensorimotor-association (S-A) axis may support age-related improvements in cognitive functioning^58^. In particular, greater segregation at each pole of the S-A axis, coupled with greater integration of middle-axis and frontoparietal systems, is thought to facilitate a balance of efficient within-system processing and cross-system communication to support developmental improvements in cognition. In line with this framework, our general exposome network consists of high within-network connectivity of the motorsensory and default mode networks and high between-network connectivity of the frontoparietal network, potentially reflecting greater segregation at sensorimotor and association poles of the S-A axis, along with the cross-system communication needed to optimize cognitive performance. Notably, prior work defining and characterizing the S-A axis has focused predominantly on cortical-cortical connectivity patterns^29,58,59^, whereas the current results also highlight an important role for subcortical and cerebellar network connectivity features. Accordingly, future work is needed to expand upon how early environments may relate to subcortical and cerebellar connectivity, and how connectivity of these networks fits within the S-A axis hierarchy to support cognitive neurodevelopment.

The present findings complement earlier work showing that the exposome is associated with spatial patterns of functional network topography, particularly for frontoparietal, default mode, and attention networks^15^. Collectively, these two sets of findings suggest convergent relationships between environmental variation and functional network topography and connectivity, particularly for the frontoparietal network. These findings are consistent with prior work suggesting that the transition to adolescence is a key period for frontoparietal network development^60^, which is thought to play a critical role in age-related improvements in cognitive functioning^58^. Notably, prior work showed that the general exposome was only weakly related to individual differences in functional network *topography* of sensory and motor networks, but the present study reveals prominent associations with the functional *connectivity* of the motorsensory network. This divergence is notable in light of research suggesting individual differences in neural topography may confound functional connectivity analyses^61^ and demonstrates that it is feasible to identify distinct associations between the exposome and functional network topography and connectivity. Together, these two sets of results suggest that the topography and connectivity of the motorsensory network may differ in developmental timing and/or susceptibility to effects of environmental exposures at the onset of adolescence.

It is interesting to note that patterns of functional connectivity predictive of general exposome scores were remarkably consistent across different brain states, including three cognitive tasks and a rest period. While there is a growing body of literature highlighting the importance of brain state for optimizing brain-behavior models^22,39–41,62^, less is known about the importance of brain state for optimizing brain-environment models that seek to characterize functional connectivity correlates of early-life environments. The current work, which leveraged an aggregate measure comprising a wide range of multivariate environmental factors, suggests that co-occurring features of childhood environments may have far-reaching impacts on youth neurodevelopment that are evident across domains and independent of ‘brain state’. Future studies may seek to further disentangle the additive and interactive effects of specific physical/chemical, psychosocial, or socioeconomic environmental factors on brain-state-specific functional connectivity patterns.

The current study leverages the large sample size and rich phenotyping of the ABCD Study dataset and applies a robust connectome-based approach incorporating stringent cross validation to elucidate patterns of functional connectivity associated with multivariate environmental exposures and youth cognition. Nonetheless, our results should be interpreted in light of several important limitations. While our models were conducted with 10-fold internal cross-validation across 100 iterations, we do not have an external validation sample. Therefore, future research is needed to externally replicate our functional connectivity findings and to assess whether the exposome network we identified is associated with cognitive functioning in independent datasets. Furthermore, the exposome was computed across all participants resulting in a lack of total independence between training and testing samples. Nonetheless, the aim of the current work is to identify patterns of connectivity associated with variation in the exposome, not to achieve true statistical prediction. Accordingly, we utilized a cross-validated CPM-style analysis approach to increase statistical rigor and adopt the term ‘connectome-based statistical inference’ to acknowledge that the current analyses do fully align with the full predictive modeling CPM framework. Additionally, the current analyses were cross-sectional and future work may examine whether the network we identified also relates to cognition at later timepoints, as well as how this network evolves across development in the context of both stable and changing environments. Moreover, given the observational design of the ABCD Study, we cannot infer causation between the exposome and identified patterns of functional connectivity. Furthermore, the general exposome does not account for all aspects of the environment that have previously been shown to be associated with brain function (e.g., racial discrimination^63,64^ and traumatic events^65,66^) and our study did not investigate non-environmental (i.e., genetic) influences on functional connectivity and cognition. Therefore, future work is needed to incorporate more facets of children’s early environments and to examine how genetic and environmental factors interact to shape cognitive neurodevelopment.

Collectively, the current results show that multidimensional features of children’s early environments are reflected in whole-brain patterns of functional connectivity at the onset of adolescence and relate to cognitive functioning across multiple key domains. These findings add to a growing literature elucidating how the environment impacts neurodevelopment and cognition among youth^67,68^. Our observation that common aspects of functional connectivity are evident across multiple key brain states (i.e., during rest, inhibitory control, reward processing, and emotional working memory task performance) highlights the far-reaching impact of the environment, as well as the potential impact of interventions geared toward improving children’s environments for fostering positive developmental trajectories^69^. Indeed, recent ABCD findings have demonstrated that associations between poverty and altered brain structure are mitigated in states that provide more generous support for low-income families^70^, highlighting the potential for policy changes to improve youth neurodevelopment. Future work is needed to investigate how the observed patterns of functional connectivity evolve across development and relate to longer-term trajectories of wellbeing.

## Data Statement

Data used in the preparation of this article were obtained from the Adolescent Brain Cognitive Development Study® (http://abcdstudy.org). Researchers with an approved NIH Brain Development Cohorts (NBDC) Data Use Certification (DUC) may obtain ABCD Study data.

## Acknowledgements

Data used in the preparation of this article were obtained from the <u>Adolescent Brain Cognitive</u> <u>Development™ (ABCD) Study</u>, held in the <u>NIH Brain Development Cohorts Data Sharing</u> <u>Platform</u>. This is a multisite, longitudinal study designed to recruit more than 10,000 children aged 9–10 and follow them over 10 years into early adulthood.

The ABCD Study® is supported by the **National Institutes of Health** and additional federal partners under award numbers:

U01DA041048, U01DA050989, U01DA051016, U01DA041022, U01DA051018, U01DA05103 7, U01DA050987, U01DA041174, U01DA041106, U01DA041117, U01DA041028, U01DA041 134, U01DA050988, U01DA051039, U01DA041156, U01DA041025, U01DA041120, U01DA0 51038, U01DA041148, U01DA041093, U01DA041089, U24DA041123, U24DA041147.

A full list of supporters is available at <u>Federal Partners – ABCD Study</u>.

ABCD Consortium investigators designed and implemented the study and/or provided data but did not necessarily participate in the analysis or writing of this report. This manuscript reflects the views of the authors and may not reflect the opinions or views of the NIH or ABCD Consortium investigators.

## Author Contributions

**Sarah D. Lichenstein**: Conceptualization, Project administration, Supervision, Writing – original draft, Writing – review and editing. **Yihe Weng**: Data curation, Formal analysis, Writing – review and editing. **Heather Robinson:** Writing – original draft, Writing – review and editing. **Lester Rodriguez:** Supervision, Visualization, Writing – review and editing. **Marzieh Babaeianjelodar**: Data curation. **Joliza Maynard:** Visualization, Writing – review and editing. **Menessa Metayer:** Visualization, Writing – review and editing. **Suhani Suneja:** Visualization, Writing – review and editing. **Corey Horien**: Data curation, Resources, Writing – review and editing. **Abigail S. Greene:** Data curation, Resources, Writing – review and editing. **Todd Constable:** Resources, Writing – review and editing. **Tyler M. Moore:** Resources, Writing – review and editing. **Ran Barzilay:** Resources, Writing – review and editing. **Sarah W. Yip**: Conceptualization, Writing – review and editing. **Arielle S. Keller:** Conceptualization, Supervision, Writing – review and editing.

## Supplementary Material

**Figure S1.**
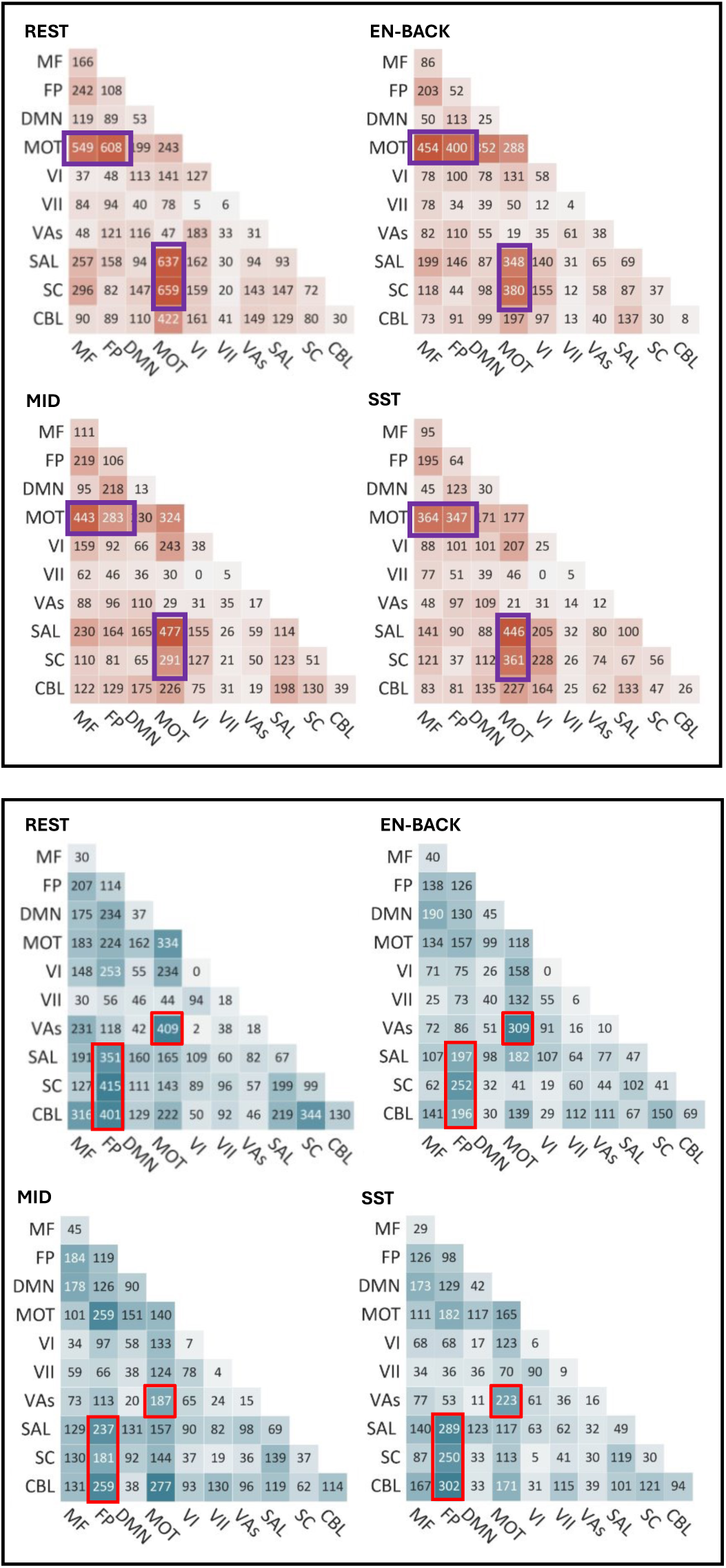
Task-specific network anatomy. Abbreviations: EN-back: emotional n-back task; MID: monetary incentive delay task; SST: stop-signal task; MF: medial frontal; FP: frontoparietal; DMN: default mode network; MS: motorsensory; VI: visual 1; VII: visual 2; VAs: visual association; SAL: salience; SC: subcortical; CBL: cerebellar.

**Figure S2.**
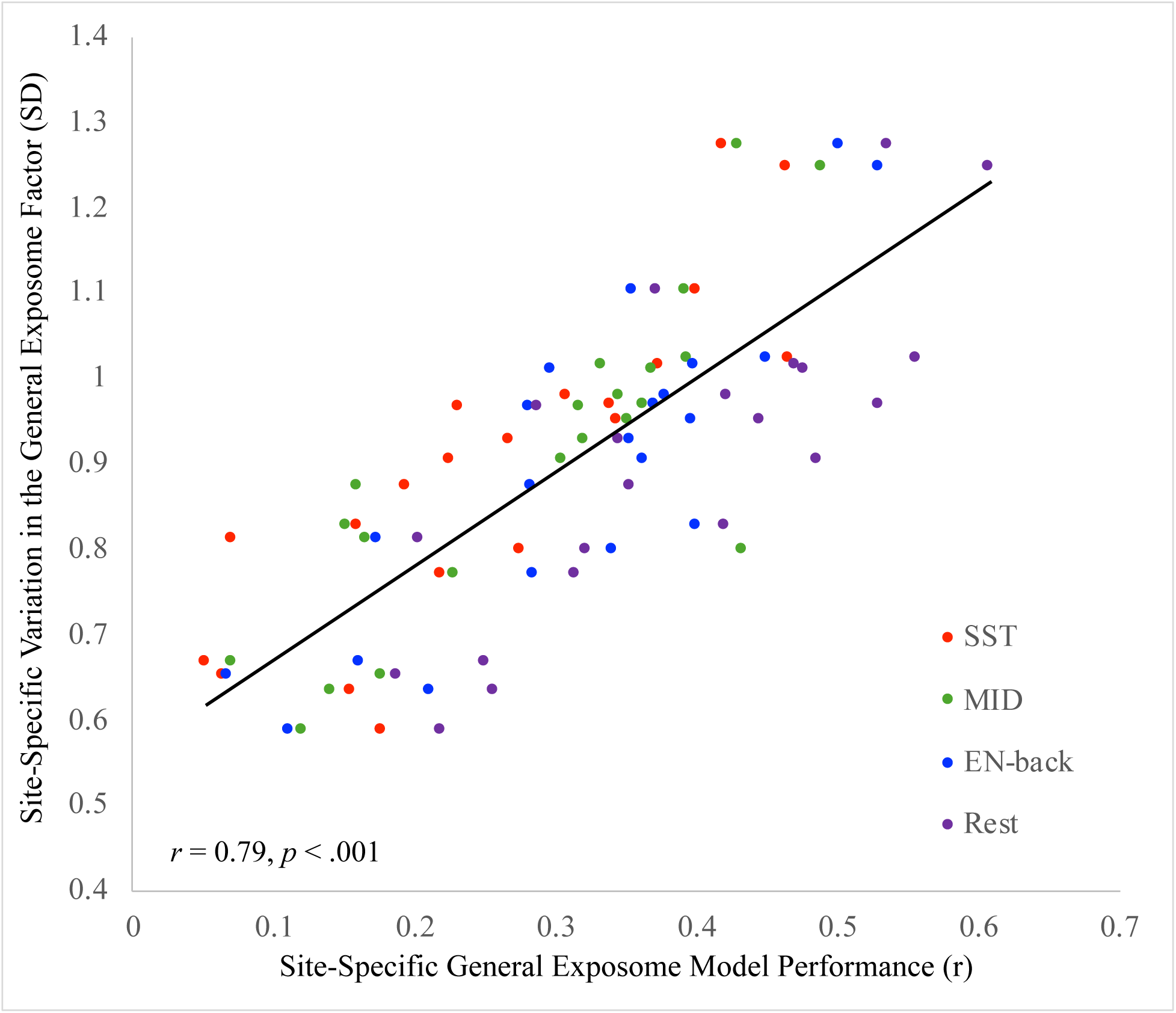
Association between site-specific model performance and exposome variation (SD)

